# Effector T cells in poorly perfused tumor regions exhibit a distinct signature of augmented IFN response and reduced PD-1 expression

**DOI:** 10.1101/2024.07.05.601540

**Authors:** Marta Riera-Borrull, Sonia Tejedor Vaquero, Víctor Cerdán Porqueras, Jose Aramburu, Cristina López-Rodríguez

## Abstract

Effector T lymphocytes are avid glucose consumers, but can function in the nutrient-poor environments of tumors. However, availability of blood-delivered nutrients throughout the tumor is not homogeneous, and how this affects effector T cells is not well known. Here we have isolated tumor-infiltrating T lymphocytes (TILs) from mouse solid tumors by their capacity to capture blood-transported probes, and compared them with glucose-restricted T cells. Glucose restriction *in vitro* arrested cell proliferation but reduced only moderately the induction of hallmark glucose-dependent cytokines interferon gamma (IFNγ) and IL-17. *In vivo*, effector TILs with reduced access to blood had characteristics of glucose-restricted cells, such as reduced expression of IFNγ and genes associated with cell proliferation. However, they expressed more CXCR3, which identifies effective antitumor T lymphocytes, showed an enhanced IFN response signature, and had reduced expression of surface PD-1. We also identified genes regulated by the enzyme ACSS2, which allows TILs to sustain gene expression in glucose-poor environments. Thus, effector T lymphocytes infiltrating tumors express different gene signatures in regions with different accessibility to blood, and can maintain specific glucose-dependent responses even in poorly perfused tumor regions. Our results can help better understand nutrient-dependent TIL heterogeneity in changing tumor microenvironments.

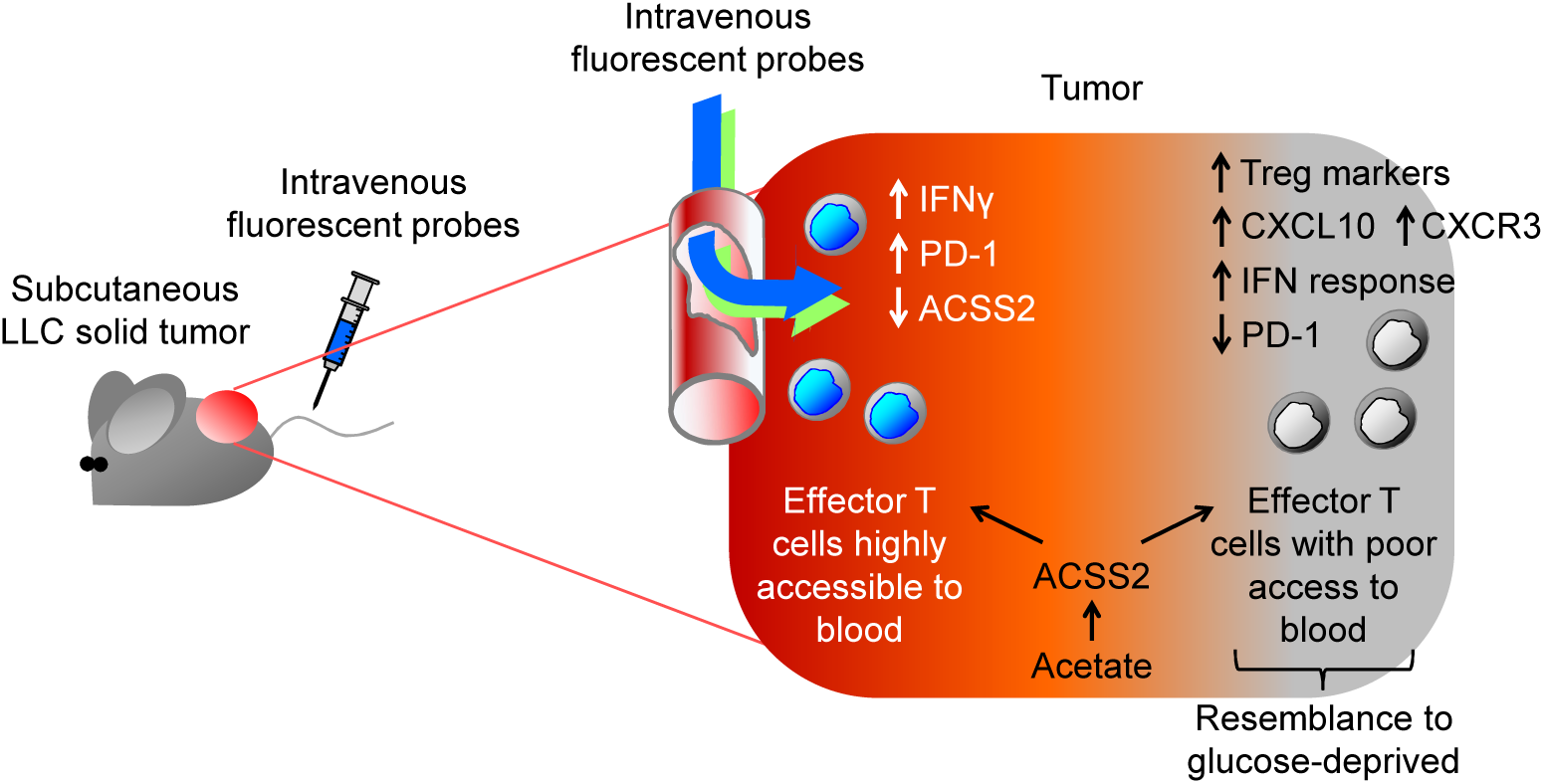

## Introduction

Activation of effector T lymphocytes is energetically demanding and requires a sufficient supply of specific nutrients. For instance, activation of naïve T lymphocytes to effector T helper (Th) 1 and Th17 cells, considered the most nutrient- and energy-hungry versions of Th cells, requires abundant glucose and glutamine (*1–3*). Accelerated glucose consumption by aerobic glycolysis is also characteristic of the reactivation of memory to effector T cells upon T cell receptor restimulation (*4*, *5*). By contrast, increased glucose processing through the Krebs cycle, and a lipid oxidation-dependent metabolism are more characteristic of regulatory T (Treg) cells, (*1*, *2*, *6–8*). Glucose is a central and very versatile nutrient for activated T lymphocytes, as it fuels rapid ATP production through glycolysis, serves as a basic building block for amino acid biosynthesis, and feeds the pentose phosphate pathway (PPP) (*9*). The PPP provides nucleotide precursors necessary to sustain T lymphocyte growth and proliferation, and is also a main source of NADPH, a major reducing agent in biosynthetic reactions and for regenerating the pool of reduced glutathione (*9*, *10*). The relevance of glucose for effector T lymphocytes is illustrated by the suppression of T cell activation by the glycolysis inhibitor 2-deoxyglucose (2DG) (*1*, *3*), and by work showing that T lymphocytes lacking the glucose transporter Glut1 fail to activate properly as effector cells and are biased toward Treg (*1*).

These observations connect with the problem that effector T lymphocytes encounter in metabolically hostile niches, such as solid tumors and infected tissues, where oxygen and glucose supply is limited (*11–18*). In contrast with lymph nodes where T lymphocyte activation occurs under optimal nutrient availability, the tumor microenvironment (TME) is one of considerable competition for nutrients between multiple cell types. This is even more critical in poorly irrigated regions, where T lymphocytes may not be able to obtain enough nutrients to sustain antitumor function. The metabolite landscape in the TME can be quite complex, with not only limited glucose availability and low oxygen levels, but also elevated levels of lactate and acetate with respect to blood or lymph nodes (*14*, *19*– *22*). In this regard, the low glucose and high lactate levels characteristic of highly glycolytic tumors can promote tumor-tolerant Treg cells instead of antitumor effector T lymphocytes (*19*, *20*). On the other hand, a glucose-poor TME might not preclude T lymphocyte responsiveness to stimulation, for instance in response to anti-checkpoint antibodies, as seen for bulk CD4 and CD8 cells as well as tumor-infiltrating Treg cells (*20*, *23*, *24*). In addition, TILs can resort to locally abundant metabolites in tumors, as shown for effector CD8 T lymphocytes metabolizing acetate to sustain interferon gamma (IFNγ) production and antitumor activity under reduced glucose availability (*21*). Altogether, these observations raise the question of which functional features can be expected from T lymphocytes in different regions within the tumor, where they face different metabolic conditions. Identifying these characteristics can help understand the heterogeneous landscape of T lymphocyte responses to tumor cells.

In this work we have analyzed glucose-dependent functions of Th17 and Th1 cells activated under very low glucose concentration, 10 to 50 times lower than in blood, which corresponds to glucose levels reported in tumors; and compared them to tumor-infiltrating effector T lymphocytes isolated from tumor regions with high or low accessibility to blood-transported probes. Previous works had analyzed the functionality and metabolic characteristics of diverse populations of tumor-infiltrating T cells, but had not identified specific features of effector TIL subsets distinguishable by their accessibility to blood. We hypothesized that TILs with reduced access to blood would be exposed to a general nutrient deficiency, and offer clues about their function compared to blood-accessible TILs. Our results reveal that T lymphocytes can achieve substantial activation and maintain energy sufficiency under very low levels of glucose. We also identify distinct gene signatures of effector TILs with very low accessibility to blood *in vivo*, and show that they express multiple glucose-dependent genes and markers associated with T cells with enhanced antitumor capacity.

## Results

### CD4 T lymphocytes can express characteristic Th1 and Th17 glucose-dependent cytokines when activated under very low glucose levels

We analyzed how glucose limitation affected the induction of primary and preactivated Th17 cells by comparing 5 mM glucose, a normal concentration in blood and lymphoid tissues, with 0.3 mM glucose, in the range (0.1 – 1 mM) measured in tumors and inflamed tissues in various pathological conditions (*13*, *15*, *25*) (Figure 1A). We initially chose Th17 cells because they are markedly glycolytic and more avid glucose consumers than other Th or Treg subsets (*2*).

**Figure 1.**
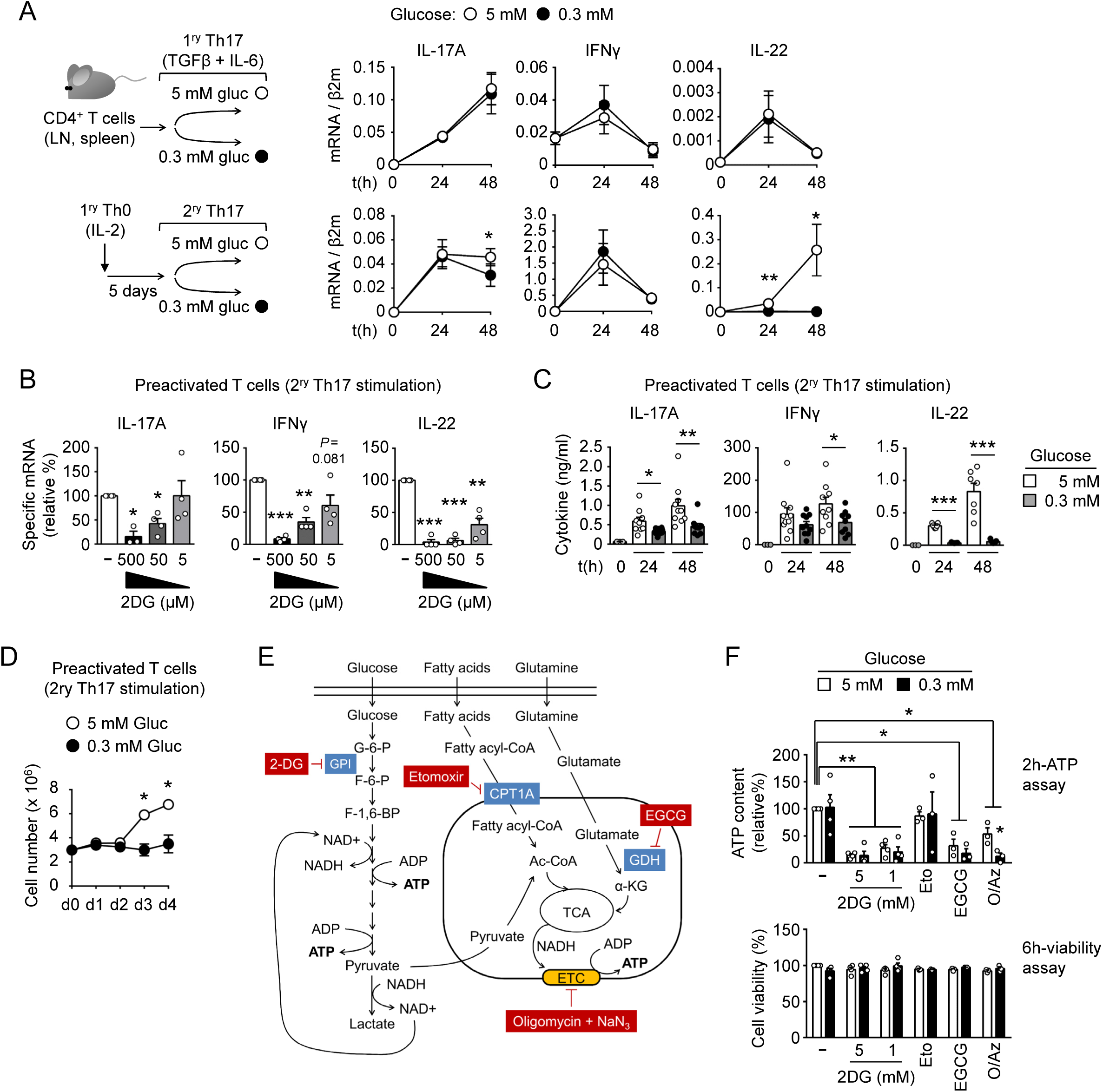
CD4 T lymphocytes can express hallmark Th1 and Th17 glucose-dependent cytokines when activated under very low glucose levels. **(A)** Upper panels show the expression of the indicated gene products in CD4 T lymphocytes freshly isolated from lymph nodes and stimulated with anti-CD3 and anti-CD28 antibodies plus TGFβ and IL-6 (primary Th17). The bottom panels show the same genes in CD4 cells that had been activated for 5 days in non-polarizing (Th0) conditions with CD3 and CD28 plus IL-2 and then restimulated as Th17 (secondary Th17) with anti-CD3 and anti-CD28 plus TGFβ and IL-6. Results show the mean ± SEM from 5 to 10 independent experiments. **(B)** Inhibition of cytokine mRNA expression in preactivated Th17 cells (48 hours) by the non-metabolizable glucose analog 2-deoxyglucose (2DG). Results show the mean ± SEM from 4 independent experiments. **(C)** Cytokine production in preactivated Th17 cells (as in the bottom panel of (A)) was measured by ELISA. Results show the mean ± SEM from 7 to 9 independent experiments. **(D)** Cell number of T lymphocytes preactivated as Th0 in 5 mM glucose and then restimulated as Th17 for 4 days in 5 or 0.3 mM glucose. Results show the mean ± SEM from 4 independent experiments. **(E)** Schematic diagram of pathways producing ATP and points of pharmacological inhibition. **(F)** Intracellular ATP content in preactivated CD4 cells restimulated as Th17 as in (A) for 24 hours and then treated with the indicated metabolic inhibitors for 2 hours (upper panel). Cell viability of parallel samples cultured for 6 hours with the inhibitors is also shown (bottom panel). Results show the mean ± SEM from 3 to 4 independent experiments. Statistical significance was assessed by an unpaired *t* test (A, C, D) or a one-sample *t* test for comparison with the reference sample (100%) in B and F. * *P* < 0.05; ** *P* < 0.01; *** *P* < 0.001. *P* values below 0.1 are also indicated.

Glucose limitation during primary Th17 stimulation did not impair the induction of IL-17A, IL-22 and IFNγ mRNA (Figure 1A). By comparison, preactivated T lymphocytes restimulated as Th17 in low glucose induced IFNγ and IL-17A mRNA comparably to cells in normal glucose, but showed a substantial decrease in IL-22 expression (Figure 1A). We confirmed that expression of these cytokines was indeed glucose-dependent as it was suppressed by the glycolysis inhibitor 2-deoxyglucose (2DG) (Figure 1B). Production of IL-17A and IFNγ protein in restimulated Th17 cells was partially inhibited under low glucose, but they still secreted substantial levels of both cytokines up to 48 hours; however, production of IL-22 was clearly impaired (Figure 1C). Finally, whereas glucose restriction did not suppress the expression of key effector cytokines in restimulated Th17 cells, it arrested their proliferation, although cultures could maintain their cellularity for at least 4 days (Figure 1D).

### Very low glucose concentrations can sustain energy production in activated CD4 T lymphocytes

We next asked whether activation of CD4 T lymphocytes in low glucose caused ATP stress or a shift in ATP production from alternative fuels such as glutamine or fatty acids. We assessed this in preactivated T lymphocytes restimulated as Th17 as in Figure 1A. Intracellular ATP content was comparable between cells cultured in 0.3 and 5 mM glucose up to 2 days (Supplemental Figure 1A). Contribution of main ATP production pathways was assessed in the last 2 hours of culture by adding different metabolic inhibitors without replacing the medium. We used 2-deoxyglucose (2DG) to block glycolysis, epigallocatechin gallate (EGCG) to inhibit conversion of glutamate to α-ketoglutarate, etomoxir for blocking the mitochondrial uptake of long-chain fatty acids, and oligomycin plus sodium azide to suppress ATP production through mitochondrial respiration and the electron transport chain (ETC) (Figure 1E). These inhibitors did not impair cell viability for at least 6 hours (Figure 1F). We observed that Th17 cells in 0.3 mM glucose obtained most of their ATP from glucose, but were more dependent on mitochondrial respiration than cells in 5 mM glucose (Figure 1F). Both were comparably dependent on glutamate to maintain their ATP levels, and did not depend on mitochondrial uptake of long-chain fatty acids (Figure 1F). We were surprised that CD4 cells activated in 0.3 mM glucose for 48 hours still had enough glucose to produce ATP. Indeed, we found that they ended up with less than 50 μM extracellular glucose by 48 hours (Supplemental Figure 1B), and even then they kept using glucose as a main nutrient for ATP production (Supplemental Figure 1C).

To find whether a similar dependence on glucose happened in other stimulatory conditions we analyzed Th1 and Th17 cells induced *ex vivo* after *in vivo* priming with anti-CD3 (*26*, *27*). *In vivo*-primed CD4 T lymphocytes reactivated as Th1 or Th17 *ex vivo* showed a variable dependence on glucose for the expression of specific cytokine mRNAs (Figure 2A). Regarding ATP, both Th1 and Th17 cells in 0.3 or 5 mM glucose were highly dependent on glycolysis and mitochondrial respiration, and largely independent of glutaminolysis and fatty acid oxidation (Figure 2B), in line with our previous results (Figure 1F). Altogether, our results suggest that CD4 T lymphocytes with different activation histories might exhibit different sensitivity to glucose limitation for the expression of specific cytokines, but still relied on glucose as a main source of ATP, even under severely limited glucose availability.

**Figure 2.**
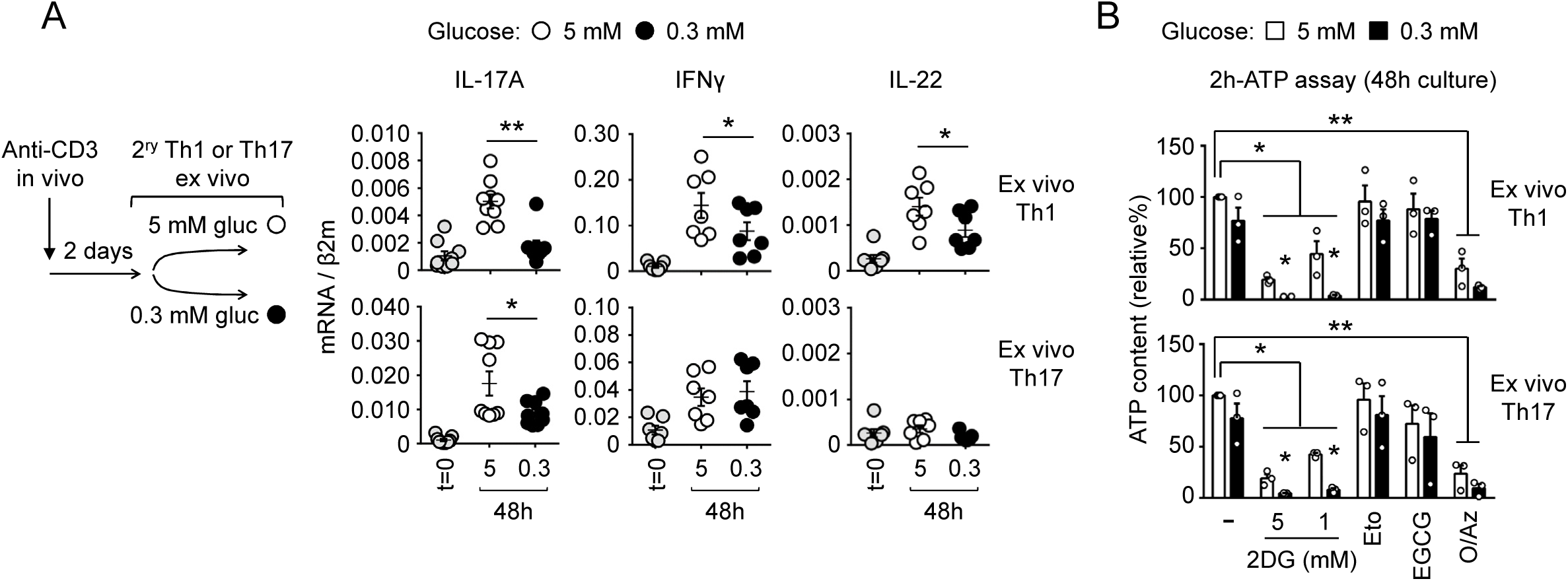
Glucose restriction has a variable impact on Th1 and Th17 cytokine expression in CD4 cells activated under different conditions. **(A)** Cytokine mRNA expression in lymph node CD4 T cells preactivated *in vivo* with injected anti-CD3 antibody and extracted from the mice 2 days later to be restimulated in culture as Th1 (with IL-12) or Th17 (TGFβ and IL-6) for 48 hours. Results show the mean ± SEM from 7 to 9 independent experiments. Statistical significance was assessed by a paired *t* test. **(B)** Intracellular ATP content in Th1 or Th17 cells stimulated as in (B) for 48 hours and then treated with the metabolic inhibitors indicated for the last 2 hours. Results show the mean ± SEM from 3 independent experiments. Statistical significance was assessed with a one-sample *t* test for comparison with the reference sample without inhibitors. * *P* < 0.05; ** *P* < 0.01.

### Delayed attenuation of mTORC1 activity under low glucose influences cytokine expression in activated CD4 T lymphocytes

Glucose sufficiency is needed for the activity of the mammalian or mechanistic target of rapamycin (mTOR), a central sensor of nutrient and energy availability that plays a major role in T lymphocyte activation (*3*, *28*). We analyzed the effect of low glucose on mTORC1 by assessing the phosphorylation of the ribosomal protein S6, a target of the mTORC1-activated kinase S6K1 (*29*, *30*). We first confirmed that mTORC1 was glucose-sensitive in our assay, as it was inhibited by 2DG (Supplemental Figure 1D). Preactivated T lymphocytes restimulated as Th17 in low glucose exhibited a mild reduction of mTORC1 activity at 24 hours, but not earlier at 6 hours (Supplemental Figure 1D). The extent of mTORC1 inhibition under low glucose was comparable to that achieved by very low concentrations of rapamycin, between 0.1 and 1 nM (Supplemental Figure 1D). These results suggested that glucose limitation during T lymphocyte activation caused a slow and progressive attenuation of mTORC1. We then assessed the effect of mTORC1 inhibition on gene expression and glucose consumption, for which we treated cells with different concentrations of rapamycin added either just before stimulation or 16 hours later to mimic a delayed inhibition of mTORC1 (Supplemental Figure 1E). Induction of IL-22 mRNA was the most mTORC1-dependent, requiring sustained mTORC1 activity; IL-17A mRNA was less mTORC1-dependent; and induction of IFNγ was mTORC1-independent, with its mRNA levels even increasing under sustained mTORC1 inhibition (Supplemental Figure 1E). Glucose consumption in these cells was also reduced by inhibiting mTORC1, but only when inhibition was done since the beginning of their stimulation (Supplemental Figure 1F).

Altogether, our results so far suggested that CD4 cells in a glucose-restricted environment are capable of maintaining an activated state for days, inducing diverse glucose-dependent cytokines, using glucose as a main fuel for ATP production, and sustaining mTORC1 activity.

### Identification of functionally distinct tumor-infiltrating T lymphocytes (TILs) *in vivo* by their uptake of the 2-deoxyglucose analog 2-NBDG

Our experiments *in vitro* showed that T lymphocytes were capable of substantial activity in glucose-restricted conditions. We then studied how T lymphocytes performed in an *in vivo* scenario of nutrient limitation. For this, we isolated effector CD4 and CD8 T lymphocytes from subcutaneously implanted Lewis lung carcinoma (LLC) tumors, based on their uptake of the 2-deoxyglucose analog 2-NBDG injected intravenously (Figure 3A and Supplemental Figure 2, A and B). When we initiated this approach, 2-NBDG uptake by T lymphocytes had been widely used as equivalent to glucose uptake (*19*, *31–36*). Recent works, though, have found that 2-NBDG and glucose are captured by T lymphocytes through different mechanisms (*24*, *37*, *38*). Nonetheless, 2-NBDG uptake is thought to correlate with the metabolic activity of T cells (*37*), and our results below indicate that TILs with poor 2-NBDG uptake resemble T cells under glucose limitation.

**Figure 3.**
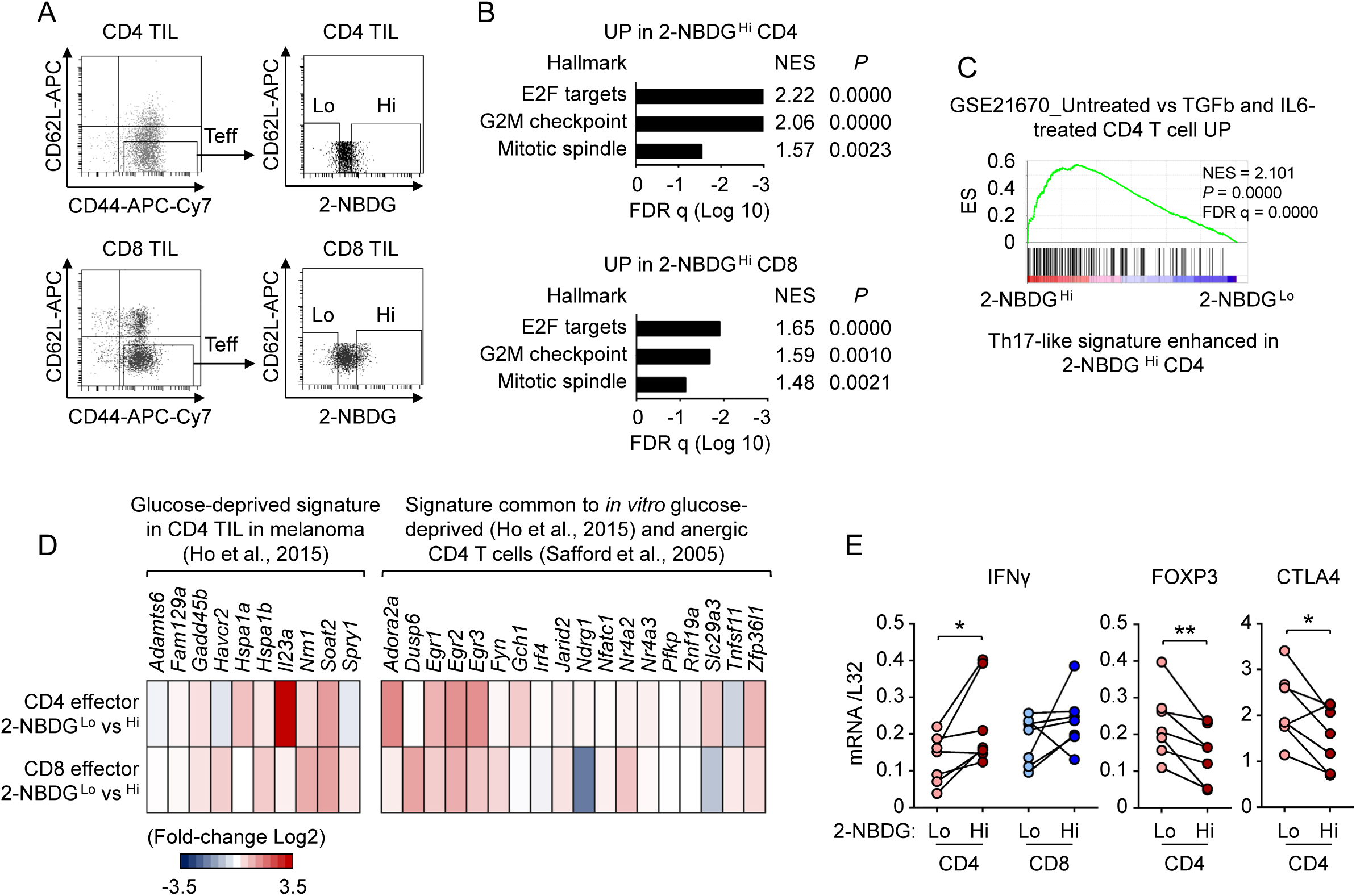
Tumor-infiltrating CD4 T lymphocytes with reduced uptake of 2-NBDG *in vivo* resemble glucose-deprived T lymphocytes and Treg cells. **(A)** Representative flow cytometry analysis of effector (CD44 ^+^, CD62L ^neg^) CD4 and CD8 TILs from LLC tumors in 2-NBDG-injected mice, gated as 2-NBDG ^Lo^ and ^Hi^ for cell sorting and RNA-seq. **(B)** GSEA of RNA-seq data shows enriched hallmarks associated with cell division in 2-NBDG ^Hi^ CD4 and CD8 TILs (NES: normalized enrichment score; *P*: *P* value; FDR: false discovery rate). **(C)** GSEA of RNA-seq data using the Immunological gene set collection database of MSigDB identifies a Th17-like signature (CD4 T lymphocytes treated with TGFβ and IL-6) enriched in 2-NBDG ^Hi^ vs 2-NBDG ^Lo^ effector CD4 TILs (NES: normalized enrichment score; *P*: *P* value; FDR: false discovery rate). **(D)** Expression of gene signatures previously associated with glucose-deprived CD4 TILs is enhanced in 2-NBDG ^Lo^ vs ^Hi^ effector CD4 and CD8 TILs. **(E)** Expression of IFNγ and the Treg-associated markers FOXP3 and CTLA4 in effector TILs sorted by their 2-NBDG uptake *in vivo*. Results in (B-D) are from one RNA-seq analysis. Results in (E) are from 2 independent experiments (4 and 3 mice respectively) analyzing effector CD4 or CD8 TILs from LLC-bearing mice injected with 2-NBDG and sorted as 2-NBDG ^Lo^ or ^Hi^. Dots are connected by lines to better visualize differences between 2-NBDG ^Lo^ vs ^Hi^ cells within each individual tumor. Statistical significance was assessed with a paired *t* test. * *P* < 0.05; ** *P* < 0.01.

We gated CD8 and CD4 TILs as effector cells (CD44 ^+^ CD62L ^neg^) and sorted them into the 15% of cells with highest and lowest 2-NBDG uptake (2-NBDG ^Hi^ and 2-NBDG ^Lo^ respectively) for RNA-seq (Figure 3A). Gene set enrichment analysis (GSEA) suggested that 2-NBDG ^Hi^ cells, both CD4 and CD8, were more engaged in cell division and proliferation than 2-NBDG ^Lo^ cells, with upregulated E2F target genes, and G2M checkpoint and mitotic spindle hallmarks (Figure 3B). We also found an enhanced Th17-like signature in 2-NBDG ^Hi^ CD4 cells, based on their similarity with T lymphocytes differentiated *in vitro* with TFGβ and IL-6 (GSE21670, (*39*)) (Figure 3C). Other characteristic Th17 markers such as IL-23R, IL-17 or IL-22 were expressed at very low levels in CD4 TILs and poorly detected in independent mRNA analyses (not shown). By contrast, 2-NBDG ^Lo^ CD4 and CD8 T lymphocytes showed generally increased expression of several markers previously associated with glucose-limited TILs, and common between anergic and glucose-restricted T lymphocytes (*25*, *40*) (Figure 3D). Additional independent experiments showed reduced IFNγ mRNA expression and increased mRNA levels of Treg markers FOXP3 and CTLA4 in 2-NBDG ^Lo^ effector CD4 TILs (Figure 3E). This result is in line with a recent report showing an enrichment in Treg-like features in 2-NBDG ^Lo^ TILs, which also exhibited low glucose uptake *ex vivo* (*19*). Altogether, these results indicate that 2-NBDG ^Lo^ effector TILs exhibit several characteristics of glucose-restricted T cells.

### Effector TILs with reduced uptake of 2-NBDG *in vivo* show an enhanced type I interferon response signature

The GSEA also uncovered the upregulation of a set of genes common to both IFN alpha and gamma responses in 2-NBDG ^Lo^ CD4 and CD8 effector TILs (Figure 4, A and B). We confirmed higher mRNA expression of IFN response genes CXCL10, CCL5, STAT1 and IFIT1 in 2-NBDG ^Lo^ cells (Figure 4C), and also found that they expressed more CXCR3 mRNA than 2-NBDG ^Hi^ cells. CXCR3 is the receptor of IFN-induced chemokines CXCL9, CXCL10 and CXCL11, and activation of CXCR3 signaling in CD8 TILs has been shown to enhance their antitumor activity (*41–43*). Additional experiments in mice lacking the type I IFN (IFN-I) receptor (IFNAR1) showed that CXCL10, IFIT1 and STAT1 were highly dependent on IFN-I signaling, whereas CXCR3 and CCL5 were less dependent on it or mostly independent (Figure 4D). These findings show that a distinct feature of 2-NBDG ^Lo^ TILs is their enhanced expression of specific IFN-I target genes.

**Figure 4.**
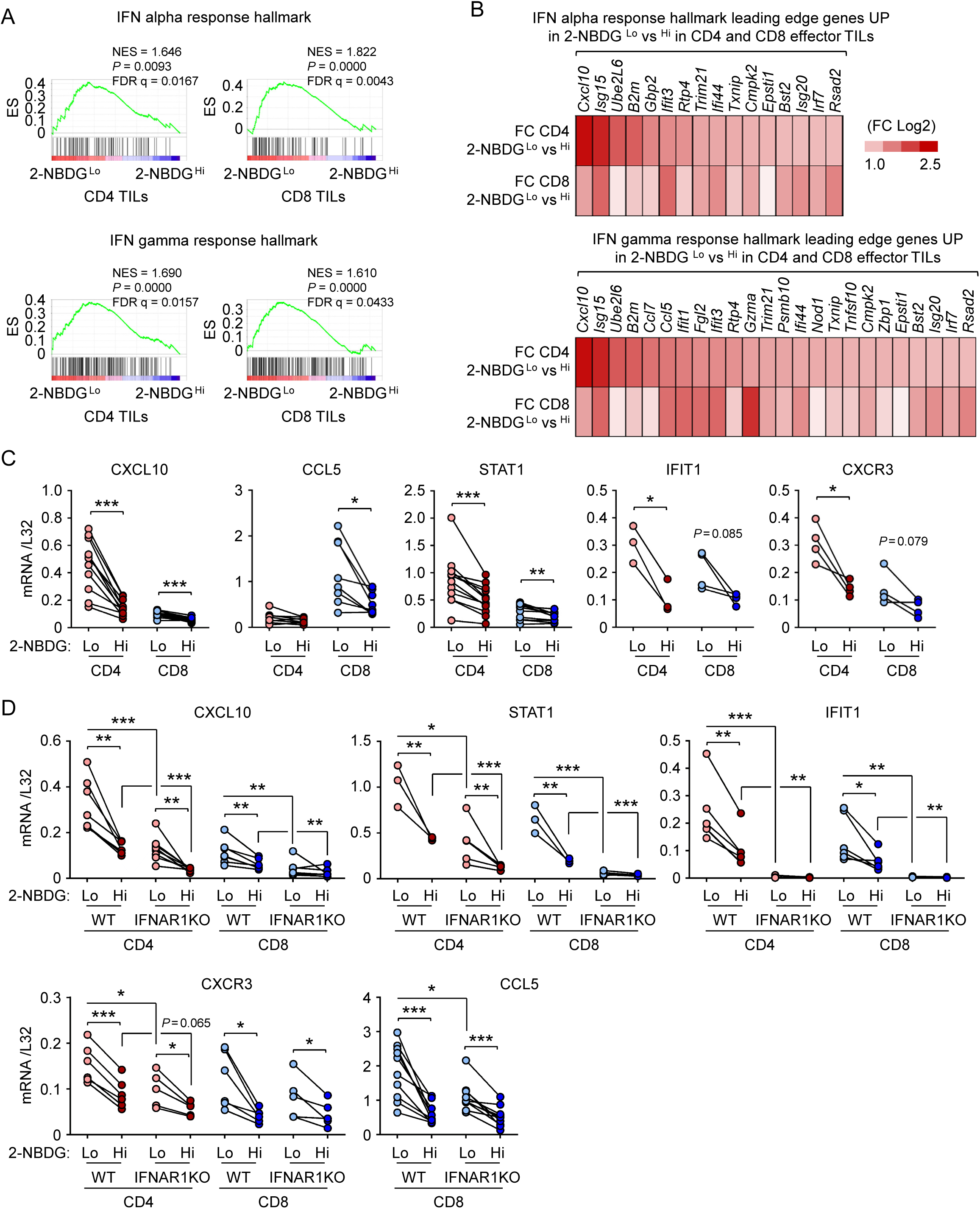
Tumor-infiltrating CD4 and CD8 T lymphocytes with reduced uptake of 2-NBDG *in vivo* exhibit an enhanced IFN response signature. **(A)** GSEA of RNA-seq data shows upregulated IFN alpha and gamma response hallmarks in 2-NBDG ^Lo^ CD4 and CD8 TILs. **(B)** IFN alpha and IFN gamma response hallmark genes with highest differential expression (fold change Log2 > 1) in both CD4 and CD8 2-NBDG ^Lo^ vs 2-NBDG ^Hi^ effector TILs in the RNA-seq data. **(C)** Expression of the indicated genes in effector CD4 and CD8 TILs sorted by their 2-NBDG uptake *in vivo*. **(D)** Expression of the indicated genes in effector CD4 and CD8 TILs sorted by their 2-NBDG uptake in vivo, in wild-type and IFNAR1-deficient mice (IFNAR1KO). Results are from 1 to 2 independent experiments. Dots are connected by lines to better visualize differences between 2-NBDG ^Lo^ vs ^Hi^ cells within each individual tumor. Statistical significance was assessed with a paired *t* test. * *P* < 0.05; ** *P* < 0.01; *** *P* < 0.001. *P* values below 0.1 are also indicated.

At this point we asked whether expression of these genes was indeed glucose-sensitive in activated T lymphocytes. *In vitro* assays showed that glucose restriction (0.3 mM) decreased the expression of IFNγ substantially in CD8 T lymphocytes, but less so in CD4 T lymphocytes (Supplemental Figure 2C). We also observed that IFNγ was unaffected by additional stimulation with IFN-I. Expression of CXCL10, IFIT1, STAT1, CXCR3, and CCL5 mRNA was also reduced in low-glucose medium (Supplemental Figure 2C). Of these, CXCL10 and IFIT1 were highly responsive to IFN-I, STAT1 less so, and CXCR3 and CCL5 were largely unresponsive (Supplemental Figure 2C), which was in line with our previous result *in vivo* in IFNAR1-deficient mice. Thus, this experiment showed that a set of genes whose expression was upregulated *in vivo* in 2-NBDG ^Lo^ TILs was glucose-dependent in cultured T cells. Since 2-NBDG ^Lo^ TILs exhibited characteristics of glucose-restricted T lymphocytes ((*19*, *25*) and Figure 3), their ability to express glucose-sensitive IFNγ and IFN response genes suggested that glucose levels in their microenvironment might be sufficient for inducing them, or that cells could use alternative metabolites.

### Effector TILs with different accessibility to blood differ in their IFN-I response and dependence on acetyl-CoA synthetase 2 for gene expression

It was possible that the reduced 2-NBDG uptake in 2-NBDG ^Lo^ TILs could be due in part to their being in tumor regions with poor access to blood. To address this, we compared 2-NBDG with an unrelated blood-transported probe, for which we coinjected intravenous 2-NBDG mixed with the DNA dye Hoechst 33342, which labels cells proportionally to their accessibility to blood vessels (*44*, *45*). We found that the bulk of CD45 ^+^ leukocytes in the tumor, as well as effector CD4 and CD8 TILs showed a good correlation between 2-NBDG and Hoechst uptake (Figure 5, A and B), indicating that 2-NBDG uptake by tumor-infiltrating leukocytes correlated in general with their accessibility to blood. This result suggested that 2-NBDG ^Lo^ and Hoechst ^Lo^ TILs were likely receiving a reduced supply of blood-delivered nutrients. As to possible mechanisms that might allow these cells to induce glucose-dependent genes, we noticed that both 2-NBDG ^Lo^ and Hoechst ^Lo^ effector TILs expressed higher levels of acetyl-CoA synthetase 2 (ACSS2) mRNA than their 2-NBDG ^Hi^ and Hoechst ^Hi^ counterparts (Figure 5C). T lymphocytes can use ACSS2 to produce acetyl-CoA, needed for gene transcription, from imported acetate abundant in the tumor microenvironment (Figure 5D), which could compensate for the reduced availability of glucose-derived acetate (*21*, *46*). Our results suggested that TILs in areas with reduced access to blood circulation could upregulate ACSS2 as an adaptive response to maintain effector functions.

**Figure 5.**
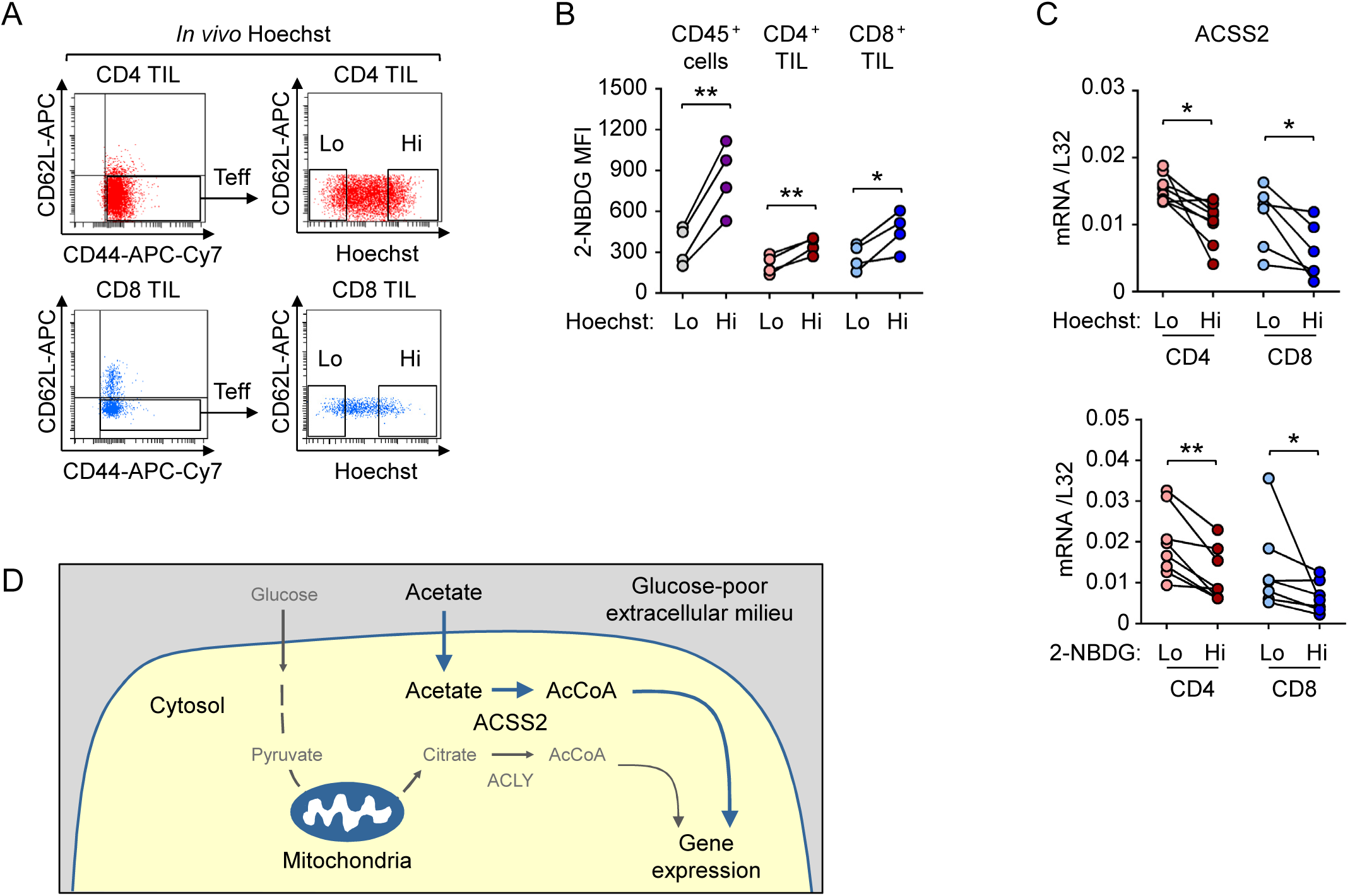
Expression of ACSS2 is elevated in tumor-infiltrating CD4 and CD8 T lymphocytes with reduced access to blood *in vivo*. **(A)** Gating strategy for sorting Hoechst ^Lo^ and ^Hi^ effector CD4 and CD8 TILs from LLC tumors in Hoechst 33342-injected mice. **(B)** Correlation between the uptake of Hoechst 33342 and 2-NBDG in immune cells (CD45 ^+^) or effector CD4 and CD8 T lymphocytes in LLC tumors, 15 minutes after intravenous injection of the probes. 2-NBDG fluorescence was quantified in the 15% lowest or highest Hoechst-stained populations. **(C)** Expression of ACSS2 mRNA in CD4 and CD8 TILs sorted by their uptake of 2-NBDG (as in Figure 3A) or Hoechst (as in (A)). Results in (C) are from 2 independent 2-NBDG and 2 independent Hoechst experiments respectively, with 3 or 4 mice each. Statistical significance was assessed with a paired *t* test. * *P* < 0.05; ** *P* < 0.01. **(D)** Diagram illustrating that T cells in a glucose-poor environment can obtain cytosolic acetyl-CoA (AcCoA) from imported extracellular acetate by using the enzyme ACSS2. Glucose normally contributes to cytosolic AcCoA when the glycolysis product pyruvate is processed through the mitochondrial Krebs cycle, from which citrate exported to the cytosol is converted to AcCoA by ATP citrate lyase (ACLY) (diagram concept adapted from *Ref 21: Qiu et al*.).

We then analyzed the contribution of ACSS2 to gene expression in effector TILs with different accessibility to blood. For this, we treated LLC tumor-bearing mice with vehicle or the ACSS2 inhibitor ACSS2i (*47*, *48*), and then isolated effector CD4 and CD8 TILs by their *in vivo* uptake of Hoechst 33342 (diagram Figure 6A). CD8 effector TILs with high Hoechst 33342 uptake showed higher expression of IFNγ, which was sensitive to ACSS2i, whereas Hoechst ^Lo^ CD8 TILs had reduced expression of IFNγ mRNA and it was ACSS2i-insensitive (Figure 6B). Hoechst ^Hi^ CD4 effector TILs did not consistently express more IFNγ than Hoechst ^Lo^ ones, but had reduced CTLA4 expression (Figure 6B). Expression of CTLA4, and also FOXP3, in CD4 TILs was partially inhibited by ACSS2i in both Hoechst ^Hi^ and ^Lo^ cells (Figure 6B). With regard to IFN response genes, Hoechst ^Lo^ CD4 TILs showed higher expression of CXCL10, IFIT1, and STAT1, as well as CXCR3 than Hoechst ^Hi^ cells. Of these genes, CXCL10 and IFIT1 were significantly dependent on ACSS2 in Hoechst ^Lo^ cells (Figure 6C). Hoechst ^Lo^ CD8 TILs also expressed more IFIT1, which was ACSS2i-sensitive, than Hoechst ^Hi^ cells (Figure 6C). STAT1, CXCR3 and CCL5 were not inhibited by ACCS2i (Figure 6C). These results show that some features of Hoechst ^Lo^ CD4 TILs, such as their elevated expression of ACSS2, CTLA4, CXCL10, IFIT1, STAT1, and CXCR3 coincided with those observed in 2-NBDG ^Lo^ CD4 TILs (Figures 3E, 4C, 5C, 6, B and C), suggesting a significant overlap between these populations. For CD8 TILs, however, there was less overlap in gene expression between 2-NBDG ^Hi^ and Hoechst ^Hi^ cells. Altogether, our results showed that ACSS2 contributed to the expression of IFNγ predominantly in CD8 TILs with higher accessibiliy to blood, to CTLA4 and FOXP3 in CD4 TILs with both high and low blood accessibility, and to CXCL10 and IFIT1 IFN-I response genes in TILs with low blood accessibility.

**Figure 6.**
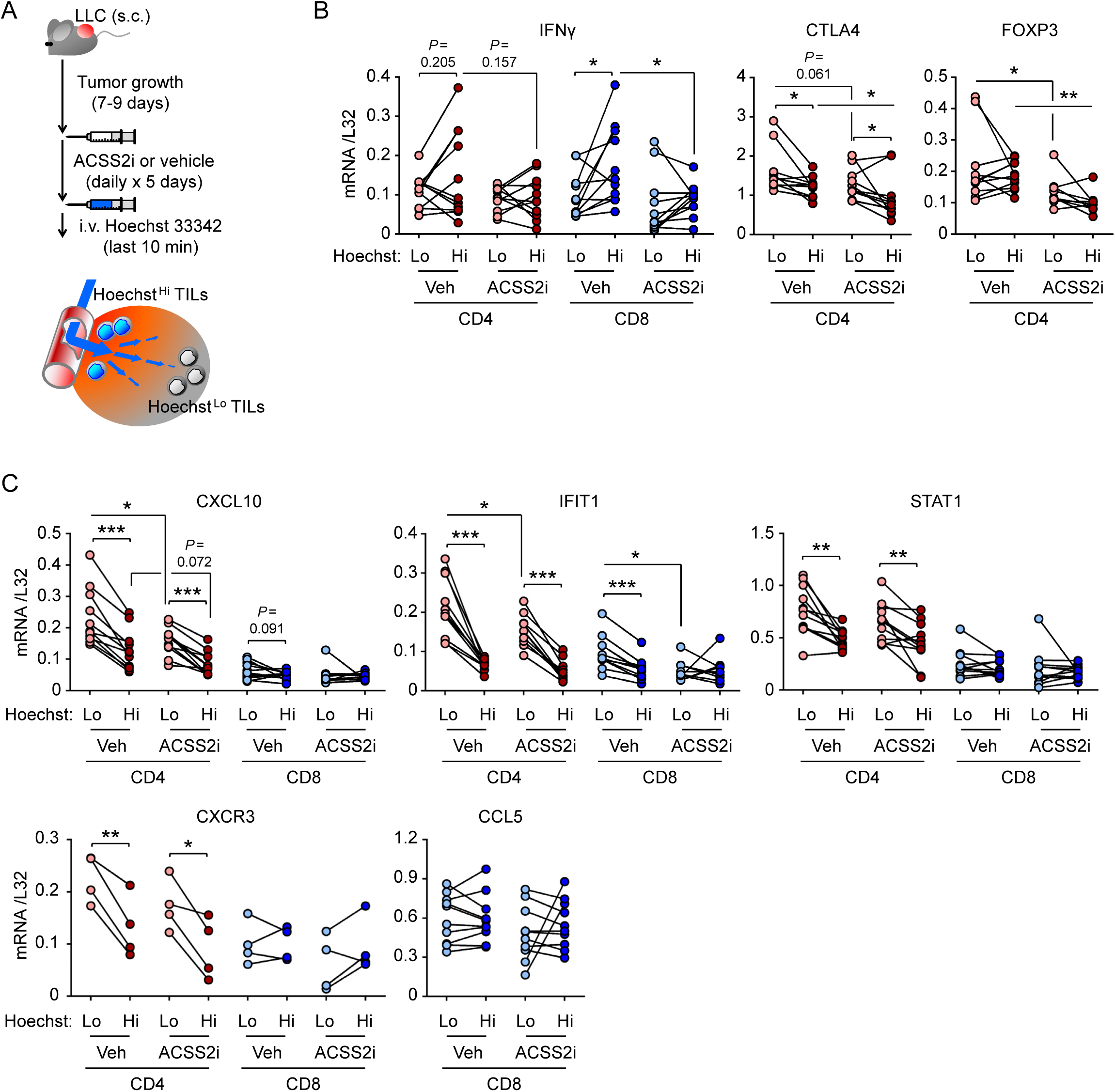
The enzyme ACSS2 regulates different genes in tumor-infiltrating CD4 and CD8 T lymphocytes with high or reduced access to blood *in vivo*. **(A)** Diagram of the experiment. **(B-C)** Expression of IFNγ, the Treg-associated markers FOXP3 and CTLA4, and IFN-response genes in effector TILs from mice left untreated or treated with the acetyl-CoA-synthetase 2 inhibitor ACSS2i, and then sorted by their uptake of Hoechst in vivo (as illustrated in Figure 5A). Results in (B-C) are from 1 to 2 independent experiments with 3 or 4 mice each. Statistical significance was assessed with a paired *t* test. * *P* < 0.05; ** *P* < 0.01; *** *P* < 0.001. *P* values below 0.1 are also indicated.

### Increased expression of CXCR3 and reduced expression of PD-1 in effector CD4 and CD8 TILs with reduced accessibility to blood

The elevated expression of CXCR3 mRNA in less blood-accessible TILs (Figures 4C and 6C) was intriguing because it has been found that CXCR3 is required for anti-PD-1-mediated T lymphocyte antitumor function (*42*, *43*). We examined the surface expression of CXCR3 in TILs and found it elevated in the Hoechst ^Lo^ and 2-NBDG ^Lo^ (less blood-accessible) fractions in both CD4 and CD8 effector TILs (Figure 7, A and B). Parallel analysis of surface PD-1 showed the opposite, with higher expression of surface PD-1 in Hoechst ^Hi^ and 2-NBDG ^Hi^ cells (Figure 7, A and B). Thus, TILs with poorer accessibility to blood displayed elevated CXCR3 and low PD-1 expression, characteristic of T lymphocytes with higher antitumor potential (*42*, *43*). Altogether, our results show that effector TILs with low accessibility to blood exhibit some characteristics of nutrient-restricted cells, but unexpectedly express an enhanced signature of glucose-sensitive IFN-I genes and a higher CXCR3 to PD-1 ratio than TILs with higher blood accessibility (diagram Figure 7C).

**Figure 7.**
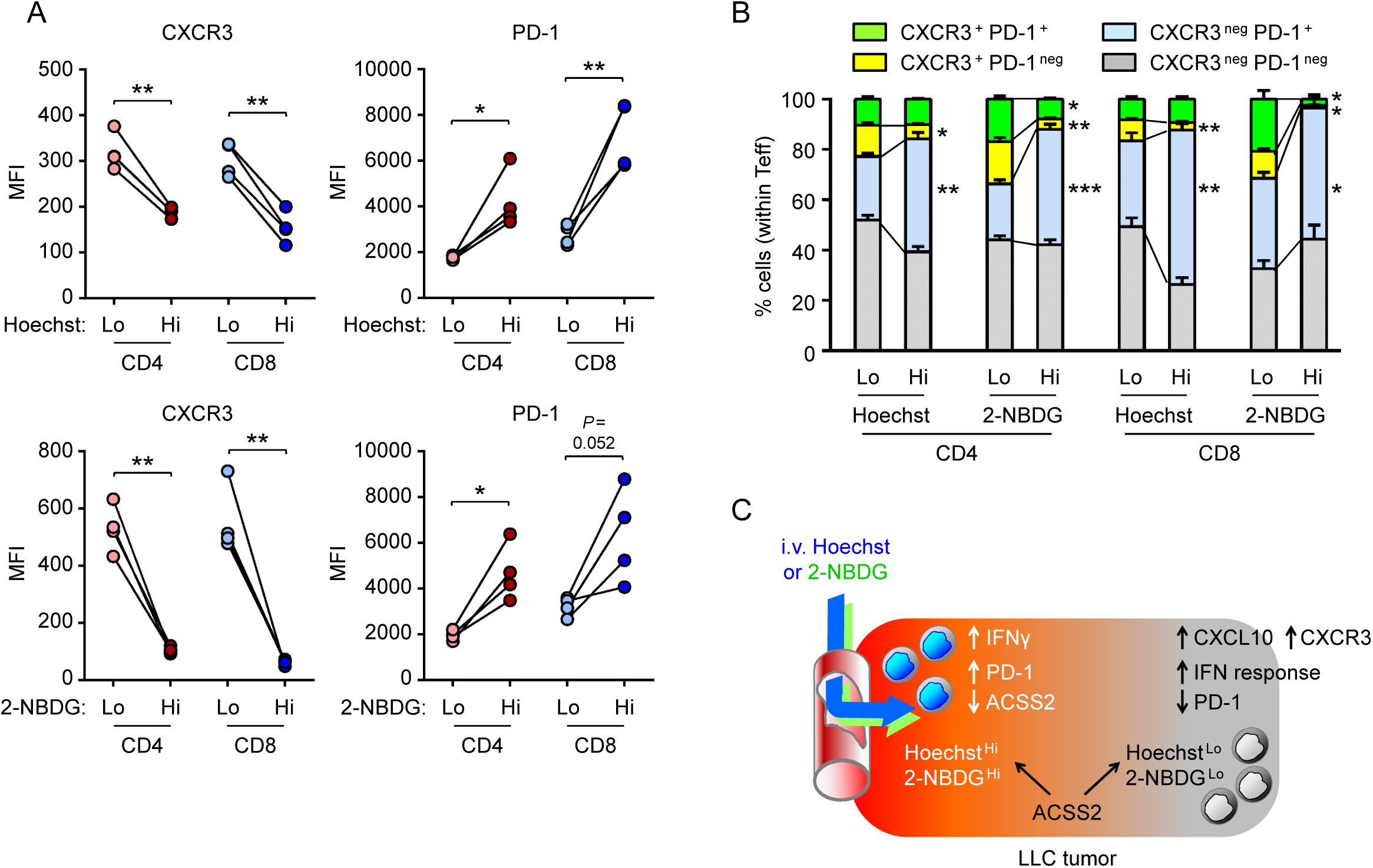
CD4 and CD8 effector TILs with reduced access to blood express more CXCR3 and less PD-1 than cells with higher blood accessibility. **(A)** Expression of CXCR3 and PD-1 surface proteins in effector TILs with low or high Hoechst uptake (upper panels) and low or high 2-NBDG uptake (lower panels), analyzed by flow cytometry, is shown as mean fluorescence intensity (MFI). Fluorescence intensity was quantified in the 15% lowest or highest Hoechst- or 2-NBDG-stained populations. **(B)** Percentage of effector CD4 and CD8 TILs double-positive for both CXCR3 and PD-1, positive for only one of them, and double-negative for both. Results are from one experiment with 4 mice. Statistical significance was assessed with a paired *t* test. * *P* < 0.05; ** *P* < 0.01; *** *P* < 0.001. *P* values below 0.1 are also indicated. **(C)** Schematic graphic model summarizing the main characteristics of TIL populations with low or high accessibility to blood-transported probes.

## Discussion

In this work we first studied the ability of activated T cells to function under substantial restriction, but not absence, of glucose, and then extended our analysis to an *in vivo* scenario, the tumor microenvironment, where effector T cells can find themselves under very limited access to blood-transported nutrients. While methods exist for measuring the capture of glucose and other nutrients *in vivo* by TILs and tumor cells using isotopic tracers, they do not allow to isolate separately cells with high or low nutrient uptake capacity. We reasoned that TILs more likely to be subjected to a general nutrient deficiency would be found in tumor regions poorly perfused by blood, and used the uptake of intravenously injected fluorescent probes by T cells as an indicator of their accessibility to blood-transported nutrients. This approach has allowed us to isolate and characterize functionally distinct populations of effector CD4 and CD8 TILs *in vivo*.

First, our *in vitro* experiments showed that effector CD4 T cells need little glucose to sustain glucose-dependent cytokine expression, energy sufficiency and mTORC1 activity, but present different glucose dependence thresholds for specific cytokines that vary with their activation history. These results are in line with recent work showing that lowering glucose down to 0.35 mM in human CD4 effector T cells severely reduces their proliferation but has a minimal impact on IFNγ production (*49*). Others have also reported that activated mouse T cells produce comparable IFNγ levels with 0.6 to 1 mM glucose as with 10 mM (*25*, *50*).

Next, *in vivo* experiments showed a significant correlation between 2-NBDG and Hoechst 33342 uptake in tumor-infiltrating leukocytes and effector CD4 and CD8 T cells, indicating that 2-NBDG uptake by TILs could parallel their general accessibility to blood-transported nutrients. Although uptake of 2-NBDG by T cells is currently considered as non-equivalent to glucose consumption (*24*, *37*, *38*), our results indicate that some characteristics of 2-NBDG ^Lo^ and Hoechst ^Lo^ effector TILs would match those of T lymphocytes with restricted access to blood-transported glucose and other nutrients. In this regard, effector T lymphocytes in poorly perfused tumor regions (2-NBDG ^Lo^ and Hoechst ^Lo^) exhibited similarities with glucose-restricted T lymphocytes in our *in vitro* assays and in earlier works (*19*, *25*, *40*), such as the downregulation of cell proliferation and cell cycle-related gene hallmarks, decreased IFNγ expression, and increased FOXP3 or CTLA4 mRNA levels. Nonetheless, it also appears that 2-NBDG ^Lo^ and Hoechst ^Lo^ effector TILs could have enough glucose in their local microenvironment to induce diverse glucose-dependent genes, including a higher expression of IFN response genes and the chemokine receptor CXCR3 than TILs with better access to blood. This possibility agrees with recent work showing that T lymphocytes in tumors might be able to capture sufficient glucose to maintain their functionality (*24*). Our results also indicate that TILs with different accessibility to blood might exhibit different metabolic adaptations, an interpretation supported by the observation that mRNA levels of the enzyme acetyl-CoA-synthetase ACSS2 were higher in 2-NBDG ^Lo^ and Hoechst ^Lo^ cells, and that TILs with low or high accessibility to blood relied differently on ACSS2 for expressing several genes. It is possible that specific metabolic adaptations in different tumor regions could confer TILs distinct functions. This idea is in line with a recent work in tumor-infiltrating γδ T cells (*51*), which used flow cytometry-based metabolic profiling (SCENITH, (*52*)) to show that IL-17-producing pro-tumor γδ T cells depend on mitochondrial respiratory metabolism, whereas IFNγ-producing antitumor cells are markedly glycolytic (*51*).

The enhanced expression of signature IFN response genes in both CD4 and CD8 effector TILs with reduced access to blood was intriguing. Elevated IFN responses are characteristic of bystander effector and memory T cells in different tissues and physiopathological situations (*53–55*). Bystander T cells in tumors may not recognize tumor-specific antigens but exhibit an activated phenotype linked to their responsiveness to locally produced cytokines, and can contribute to antitumor immunity (*56*). As IFN-I can strengthen antitumor T lymphocyte activity (*57–59*), a heightened IFN-I response in TILs with limited access to blood could help them retain antitumor capability.

Effector CD4 and CD8 TILs with reduced blood accessibility also showed enhanced expression of the chemokine receptor CXCR3. CXCR3 ^+^ CD4 TILs can have Treg characteristics (*60*), whereas CXCR3 ^+^ CD8 TILs are capable of strong antitumor activity in response to anti-checkpoint immunotherapy (*42*, *43*). CXCR3 has been recently shown to be highly expressed in tumor-infiltrating CD44 ^+^ CD4 Treg cells. These cells interact with CXCL9-expressing type 1 dendritic cells (DC1), preventing them from stimulating antitumor cytotoxic CD8 cells (*60*). Also, TILs in poorly perfused areas are likely to coexist with tumor-tolerant macrophages, which are more abundant in tumor regions poorly accessible to Hoechst 33342 (*44*), and this could reinforce Treg influence locally. On the other hand, the elevated expression of the IFN-inducible chemokine CXCL10 in Hoechst ^Lo^ and 2-NBDG ^Lo^ CD4 TILs could provide a stimulatory feedback loop for CXCR3 ^+^ effector CD8 cytotoxic TILs, since CXCR3 not only drives chemotaxis but also stimulates their antitumor capacity in response to anti-PD-1 immunotherapy (*42*, *43*). Thus, changes in expression of CXCL10 and CXCR3 in CD4 and CD8 TILs depending on their location within the tumor could influence a local balance between tumor-tolerant and antitumor T cell activities.

Finally, TILs with low accessibility to blood expressed less surface PD-1 than highly accessible cells. These results suggest that effector TILs with lower accessibility to blood might respond to anti-checkpoint immunotherapy differently from more blood-accessible TILs, not only because they express less PD-1 but also because they might be less accessible to blood-transported antibodies (*61*). Also, differences in the proportion of effector and Treg cells between tumor regions with better or poorer access to blood could affect the outcome of anti-PD-1 treatment, as suggested by work showing that the suppressive capacity of human Tregs in glucose-low, lactate-rich media is enhanced by anti-PD-1 treatment (*23*). In this regard, it remains to be determined which type of CXCR3 ^+^ T lymphocytes are enriched in poorly accessible tumor regions, whether Treg or antitumor cells; and how TILs in highly and poorly blood-accessible tumor regions would respond to anti-PD-1 immunotherapy.

In sum, our work has uncovered different functional and metabolic characteristics of effector tumor-infiltrating T lymphocytes with better or poorer access to blood. Our results suggest that these differences could influence the outcome of tumor-immune system interactions and T lymphocyte responses to immunotherapy approaches.

## Materials and Methods

### Mice

C57Bl/6 wild-type mice were bred and housed in specific pathogen-free conditions at the animal facility of Parc de Recerca Biomèdica de Barcelona (PRBB). *Ifnar1* ^−/−^ mice, described in (*62*), were kept in a C57BL/6 background and were obtained from Manuel Rebelo at the Rodent Facility of the Gulbenkian Institute (Lisbon, Portugal). Male and female animals were used, and similar findings were obtained for both sexes.

### T lymphocyte isolation and culture

Mouse CD4^+^ T lymphocytes were obtained from spleen or lymph nodes with the MagnisortTM Mouse CD4 T lymphocyte Enrichment Kit (eBioscience, catalog: 8804-6821-74) according to the manufacturer’s instructions. Cells were activated as Th0 with hamster anti-mouse CD3 (1 μg/10^6^ cells) plus hamster anti-mouse CD28 (1 μg/10^6^ cells) antibodies (BD Biosciences, catalog: 553058 and 553295 respectively) in plates coated with goat anti-hamster IgG (9.5 μg/cm^2^). Cells were cultured at 1 to 1.5 x 10^6^ cells/ml in DMEM glucose-free medium (Gibco, catalog: 11966025) supplemented with 10% fetal bovine serum (FBS, Life Technologies, catalog: 10270-106), non-essential amino acids (Life technologies, catalog: 11140-035), 2 mM L-glutamine (Life technologies, catalog: 25030024), 50 μM β-mercaptoethanol (Gibco, catalog: 31350-010), 10 mM Hepes (Life technologies, catalog: 15630-056) and penicillin and streptomycin (Gibco, catalog 15140122). Glucose in this culture medium was provided only by the 10% FBS, and was measured to be 0.3 mM. T lymphocyte cultures were done in medium with either no additional glucose supplementation (0.3 mM glucose), or supplemented with glucose (Life Technologies, catalog: A2494001) up to 5 or 15 mM as indicated in the respective figures. Th0 cultures were supplemented with 5 ng/ml recombinant human IL-2 (Proleukin; Chiron, formerly Eurocetus, Emeryville, CA, USA). After 48 hours and depending on visual estimation of cell density, cells were split 1/2 or 1/3 in fresh medium supplemented with 5 ng/ml IL-2. For expansion of preactivated CD4 T lymphocytes under Th17 polarization, Th0 cells were restimulated at 10^6^ cells/ml with anti-CD3 and anti-CD28 antibodies as above, plus 10 ng/ml IL-6 (ImmunoTools, catalog: 12340063) and 2.5 ng/ml TGFβ (PeproTech, catalog: 100-21) for up to 4 days. For Th1 polarization T lymphocytes were cultured with 5 ng/ml of recombinant murine IL-12 (Peprotech, catalog 210-12). For stimulation of preactivated T lymphocytes with type I IFN, T lymphocytes (both CD4 and CD8) isolated with Dynabeads flowcomp mouse pan T (Thermofisher, catalog: 11465D) from lymph nodes were first activated as Th0 with anti-CD3 and anti-CD28 plus IL-2 as indicated for CD4 cells above. After 5 days, T lymphocytes were harvested, adjusted at 10^6^ cells/ml in fresh medium (15 mM or 0.3 mM glucose) and restimulated with anti-CD3 and anti-CD28 antibodies plus IL-2 (5 ng/ml) for 24 hours without or with additional stimulation with a cocktail of 600 U/ml IFNα4 (recombinant mouse IFN alpha 4, R&D Systems, catalog: 12115-1) and 2.5 ng/ml IFNβ1 (recombinant mouse IFN-β1 (carrier-free), Biolegend, catalog: 581302). Cells were then isolated as CD4 or CD8 respectively with immunomagnetic beads (Dynabeads Mouse CD4 ^+^, Invitrogen, catalog: 11445D; or Dynabeads Sheep anti-Rat IgG, Invitrogen, catalog: 11035 coated with anti-mouse CD8 rat hybridoma 53-6.7)

### Gene expression analysis

Total RNA from T lymphocytes was isolated using the High Pure RNA isolation kit (Roche, catalog 11828665001), quantified in a NanoDrop (ND-1000) spectrophotometer and 100 ng to 300 ng of total RNA was retrotranscribed to cDNA using the First Strand cDNA synthesis kit with random hexamers (Roche, catalog 04 897 030 001), or the High-Capacity cDNA Reverse Transcription Kit with RNAse inhibitor from Applied Biosystems (catalog: 4374967). Gene expression was analyzed by real-time quantitative PCR (RT-qPCR) using LightCycler 480 SYBR Green I Master Mix (Roche, catalog 4729749001) and the LightCycler 480 Real-Time PCR System (Roche) according to the manufacturer’s instructions. Primers used for RT-qPCR are listed in Table S1.

### Enzyme-linked immunosorbent assay (ELISA)

Cell-free supernatants from cultures of isolated CD4^+^ T lymphocytes (1 x 10^6^ cells/ml) were harvested and stored at -80°C. Detection of IL-17A, IL-22 and IFNγ was done in duplicate in 96 well-plates (Costar, catalog: 3590) with the Mouse ELISA Ready-Set GO® system (eBioscience, catalog: 88-7371-88, 88-7422-88 and 88-7314-88 respectively) following the manufacturer’s instructions.

### ATP measurement

At different times upon culture in 5 or 0.3 mM glucose medium, T lymphocytes were adjusted at 50.000 cells/ml in their own culture medium and 0.5 ml of the cell suspension were transferred to 1.5 ml microcentrifuge tubes and incubated at 37°C for 2 hours in the absence or presence of metabolic inhibitors: 50 to 1 mM 2-deoxyglucose (2-DG) (Sigma, catalog: D8375-5G), 200 µM etomoxir (Sigma, catalog: E1905-25), 50 µM epigallocatechin gallate (EGCG) (Sigma, catalog: E4143), 0.1 µg/ml oligomycin and 20 mM sodium azide. Cells were harvested, washed with a solution containing 150 mM NaCl and 50 mM Tris pH 8, resuspended in 100 µl of wash solution plus 0.2% Triton and frozen at -80°C. ATP concentration in cell lysates was measured in a FB 12 Luminometer (Berthold detection systems) using the ATP Bioluminiscence Assay Kit CLS II (Roche).

### Glucose measurement

Cell-free supernatants from cultures of isolated CD4+ T lymphocytes (1 x 10^6^ cells/ml) were harvested and stored at -80°C. Glucose measurements were done in duplicate in 96 well-plates (Costar, catalog: 3590) using the Glucose Assay kit I (Eton Bioscience, catalog 12000310) according to the manufacturer’s instructions.

### *In vivo* anti-CD3 injection and *ex vivo* restimulation

8-12 week-old C57BL/6 female mice were injected intraperitoneally with anti-CD3 1 mg/kg (clone 145-2C11, DB Bioscience, catalog: 553058) to induce systemic inflammation and T lymphocyte activation (*26*, *27*). After 48 h, CD4^+^ T lymphocytes were isolated from peripheral lymph nodes, and restimulated *ex vivo* with anti-CD3 plus anti-CD28 (1 µg/10^6^ cells) in plates coated with goat anti-hamster antibody (Goat anti-hamster IgG affinity-purified antibody, MP Biomedicals, catalog: 856984), with the glucose concentration and polarizing conditions indicated in Figure 2A.

### Intracellular staining for phospho-S6 and flow cytometry

T lymphocytes were fixed with a fixation/permeabilization solution (eBioscience, catalog: 00-5521) during 40 minutes to 1 hour at 10^5^ cells/100 µl. They were then washed twice with permeabilization buffer (Permwash, eBioscience, catalog: 00-8333-56) and permeabilized for 20 minutes in the same solution. Anti-phospho-S6 (S235/236) antibody (Cell Signaling Technology, catalog: 2211S) was then added at a 1:100 dilution and incubated for 2 hours at 4°C. After washing cells twice with permwash solution, cells were incubated with a secondary FITC-labeled antibody (donkey anti-rabbit) for 20 minutes at 4°C. Cells were then washed twice and resuspended in PBS and analyzed with an LSR II flow cytometer. Data analysis was done using the FlowJo software (TreeStar).

### Lewis lung carcinoma (LLC) tumors, *in vivo* labeling with 2-NBDG and Hoechst 33342

LLC cells derived from C57BL/6 mice were kindly provided by Dr. I. Melero (Center for Applied Medical Research, Pamplona, Spain). LLC cells were grown in complete medium and maintained at subconfluency by passing them with gentle pipetting. LLC were routinely tested and confirmed to be mycoplasma-free. For solid tumor development, 5 × 10^5^ LLC cells per mouse were injected subcutaneously in the right back flank of 8–12 week old C57BL/6 mice. Male and female mice were used. Starting at day 7 post-inoculation, tumors were measured every other day using a caliper and the tumor volume was calculated using the formula L × W^2^ × 0.52, where L = maximal length and W = maximal width. Mice were sacrificed between day 12 to 15 after tumor inoculation, when tumors reached approximately 500 mm^3^ on average.

*In vivo* labeling of tumor-infiltrating T lymphocytes (TILs) with 2-(N-(7-nitrobenz-2-oxa-1,3-diazol-4-yl) amino)-2-deoxyglucose (2-NBDG, Life Technologies, catalog: N13195) was done by intravenous (retroorbital) injection of 2-NBDG (20 mg/kg of mouse body weight; 3.33 mg/ml in PBS). 2-NBDG was allowed to circulate for 15 min before euthanizing the mice and excising the tumors to isolate TILs. Labeling with Hoechst 33342 (Thermofisher, catalog: H3570; 12 mg/kg of mouse body weight; 3.33 mg/ml in PBS) was done in the same way.

Excised LLC tumors were minced in 1.5 ml tubes containing 0.5 ml complete DMEM without glutamine and β-mercaptoethanol. The content was transferred to a 50 ml tube containing 2.5 ml of the same medium to which 0.5 mg/ml of collagenase A (Roche, 10103578001) and 0.01% DNase I (Sigma-Aldrich, D4263-5VL) were added. Tumors were digested for 40 to 45 min at 37°C under continuous rotation, filtered through 70 μm cell strainers to remove undigested fragments, and transferred to 15 ml tubes. Cells were then centrifuged (330 x g for 5 min), incubated with 1 ml erythrocyte lysis buffer (Biolegend, 420301) for 7 min at 4°C, then washed with 10 ml PSA buffer (PBS with 10% FBS and 0.1% sodium azide) and centrifuged again. Tumor pellets were resuspended in PSA to obtain single-cell tumor suspensions for flow cytometry staining or sorting.

### Flow cytometry and cell sorting of tumor-infiltrating T lymphocytes

Samples were analyzed with an LSRII cytometer (BD biosciences) equipped with 355, 405, 488, and 633 nm lasers, or an LSR Fortessa (BD biosciences) equipped with 405, 488, 561, and 633 nm lasers. For cell sorting, cell suspensions were filtered through a 35-μm nylon mesh (Falcon, Cat# 352235) and isolated with a BD FACSAria II (BD biosciences) cell sorter equipped with 355, 405, 488, 561, and 633 nm lasers. Experiments were analyzed with FACSDiva versions 6.2 and 8 software (BD biosciences, http://www.bdbiosciences.com/us/instruments/research/software/ flow-cytometry-acquisition/bd-facsdiva-software/).

Effector CD4 and CD8 TILs were identified as CD44 ^+^ CD62L ^neg^ within alive T lymphocytes cells gated as CD45.2 ^+^, CD90.2 ^+^ (Thy1.2), and CD4 ^+^ or CD8 ^+^ respectively. The antibodies used were CD45.2 PE-Dazzle 594, Biolegend, catalog: 109846; CD90.2-PE-Cy7, Biolegend, catalog 140310; CD4-PE-Cy5, eBioscience, catalog: 15-0042-83; CD8-PE, Biolegend, catalog: 100708; CD44-APC-Cy7, Biolegend, catalog: 103028; CD62L-APC, eBioscience, catalog: 17-0621-83; CD279 (PD-1) Brilliant Violet 605, Biolegend, catalog: 135220; and CXCR3 Alexa Fluor 700, R&D Systems, catalog: FAB1685N. Cell sorting of 2-NBDG and Hoechst 33342 high or low populations was done on the 15% of CD4 and CD8 effector cells with the highest or lowest staining with each respective dye.

### RNA sequencing and bioinformatics analysis

Effector CD4 or CD8 TILs from LLC tumor-bearing mice were sorted as 2-NBDG ^Hi^ or 2-NBDG ^Lo^ from 4 independent mice. The 15% of effector TILs with the highest 2-NBDG staining were sorted as 2-NBDG ^Hi^ cells, and the lowest 15% were sorted as 2-NBDG ^Lo^. For each population, a pool of 2000 cells (500 cells per mouse) was done. RNA was extracted with the RNeasy Microkit (Qiagen, Catalog: 74004), and cDNA libraries were prepared from poly A RNA (mRNA) using the Smart-Seq2 single-cell protocol (*63*) with some modifications. Briefly, reverse transcription was done with SuperScript™ II reverse transcriptase (Invitrogen, Catalog: 18064014), oligo-dT30VN (1 µM; 5′-AAGCAGTGGTATCAACGCAGAGTACT30VN-3′), template-switching oligonucleotides (1 µM) and betaine (1 M). Template switching was done using a locked nucleic acid (LNA) without the purification step before preamplification PCR to obtain an increased cDNA yield. cDNA concentration was measured with the Qubit dsDNA High Sensitivity assay (Invitrogen, Catalog: Q32851) and analyzed with Agilent Bioanalyzer or Fragment analyzer High Sensitivity assay (Agilent, Catalog: 5067-4626 or DNF-474) to check the size distribution profile. cDNA libraries were prepared using NEBNext® Ultra DNA Library Prep for Illumina® kit (NEB, catalog: E7370) according to the manufacturer’s protocol. 5 ng of cDNA were fragmented at a range size of 200-500bp using Covaris S2, then subjected to end repair and addition of “A” bases to 3′ ends, ligation of adapters and USER excision. All purification steps were performed using AgenCourt AMPure XP beads (Beckman Coulter, catalog: A63882,). Library amplification was performed by PCR using NEBNext® Multiplex Oligos for Illumina (Index Primers Set 1, Catalog E7335), (Index Primers Set 2, Catalog: E7500), (Index Primers Set 3, Catalog E7710) or/and (Index Primers Set 4, Catalog: E7730). Final libraries were analyzed with Agilent Bioanalyzer or Fragment analyzer High Sensitivity assay to estimate the quantity and check size distribution, and were then quantified by qPCR using the KAPA Library Quantification Kit (KapaBiosystems, Catalog: KK4835,) prior to amplification with Illumina’s cBot. Libraries were sequenced 1 x 50 + 8 bp on Illumina’s HiSeq2500.

The quality of the raw data was checked using FastQC (version 0.11.5) (*64*) and trimmed using Skewer (version 0.2.2) (*65*). The percentage of reads mapping to ribosomal RNA was assessed using riboPicker (version 0.4.3) (*66*). RNA-seq raw reads (31 to 36 x 10^6^ reads/sample) were mapped to the *Mus musculus* reference genome (Gencode M21 release; mm10; files GRCm38.primary_assembly.genome.fa.gz and gencode.vM21.annotation.gtf) and counted at the gene level using STAR (version 2.5.3a) (*67*) and parameters ’--outSAMunmapped None --outSAMtype BAM SortedByCoordinate --quantMode GeneCounts’. As the library preparation protocol was reverse stranded, the 4^th^ column of the STAR gene counts output of each sample was taken as input for DESeq2. Qualimap (version 2.2.1) (*68*) was used to check the quality of the aligned reads. Pre-Ranked Gene Set Enrichment Analysis (GSEA) (*69*) was used to obtain the functional annotation of gene lists, using the Molecular Signatures Database (MSigDB) collections Hallmark and Immunological signature gene sets. Genes with less than 30 reads across all samples were filtered out before processing the data. The accession number for the RNA-seq datasets is GSE250248.

### *In vivo* treatment with ACSS2i

Mice bearing LLC subcutaneous tumors were injected intraperitoneally with 300 µl of ACSS2i (25 mg/kg body weight) or vehicle (10% DMSO + 40% PEG 300 + 5% Tween 80 + 45% saline), beginning at day 7-9 after tumor inoculation and daily during 5 days (*47*, *48*). Mice were euthanized 16 hours after the last administration of ACSS2i/vehicle. 10 minutes before euthanasia mice were injected intravenously (retroorbital) with Hoechst 33342 to label TILs with higher or lower accessibility to blood.

### Statistical analysis

Statistical analyses were done with GraphPad Prism 6. Significance of the differences between sets of experimental data was determined with an unpaired Student’s *t* test, with Welch’s correction whenever variances were significantly different between the compared groups. A one-sample *t* test was used when samples were compared to a reference control sample. Paired *t* tests were used to compare 2-NBDG-high versus -low and Hoechst 33342-high versus -low cells, sorted from the same tumor, and when comparing different conditions of the same culture tested simultaneously.

### Study approval

Animal handling and experiments were in accordance with approved protocols by the Parc de Recerca Biomèdica de Barcelona Animal Care and Use Ethics Committee and carried out in accordance with the Declaration of Helsinki and the European Communities Council Directive (86/069/EEC).

## Supporting information

Riera-Borrull et al Supplemental material

## Data availability

RNA sequencing datasets of CD4 and CD8 effector T cells isolated from tumors have been uploaded to GEO under the accession number GSE250248. Values for all data points in graphs are reported in the Supporting Data Values file.

## AUTHOR CONTRIBUTIONS

Conceptualization: JA, CLR; Methodology and analysis: MRB, STV, VCP, JA, CLR; Experiments were performed by: MRB, STV, VCP; Supervision: JA, CLR; Writing original draft: JA; Review: VCP, STV, MRB, JA, CLR: Editing: JA, CLR; Funding acquisition: JA, CLR.

## ACKNOWLEDGMENTS

We would like to thank Òscar Fornas, Erica Ramírez, Eva Julià and Alexandre Bote of the Universitat Pompeu Fabra and Center for Genomic Regulation (UPF/CRG) Flow Cytometry Unit for excellent guidance with flow cytometry. We also thank Jochen Hecht, Irene Navarrete (CRG) and the CRG Genomics Unit for advice with and performing the RNA-sequencing; and Julia Ponomarenko and Sarah Bonnin (CRG) for bioinformatics analysis of RNA-sequencing data.

## Funding

CLR and JA were funded by: grants RTI2018-095902-B-I00 and PID2021-128721OB-I00, funded by *Agencia Estatal de Investigación* MICIU/AEI/ 10.13039/501100011033 and by European Fund for Regional Development “ERDF/EU *A way of making Europe*”; and Worldwide Cancer Research: WWCR, 20-0144. Additional funding to CLR was from ICREA Acadèmia program (2014 and 2022 awards) of *Institució Catalana de Recerca i Estudis Avançats* (ICREA, *Generalitat de Catalunya*). STV was funded by a predoctoral fellowship of the Spanish Ministry of Economy and Competitiveness: BES-2013-062670. VCP was funded by a predoctoral fellowship by the program “*Formación de Profesorado Universitario*” of the Spanish Ministry of Universities: FPU19-04119. Additional funding to CLR and JA laboratory was from the *Departament de Recerca i Universitats*, *Generalitat de Catalunya* 2021SGR00683; and the Spanish Ministry of Science Innovation through the “*Unidad de Excelencia María de Maeztu*” program funded by the *Agencia Estatal de Investigación*-AEI: CEX2018-000792-M.

Address correspondence to: Cristina López-Rodríguez, Department of Medicine and Life Sciences, Universitat Pompeu Fabra, Carrer Dr Aiguader 88, 08003 Barcelona, Spain. Telephone number: +34 93 3160810. Or to: Jose Aramburu, Department of Medicine and Life Sciences, Universitat Pompeu Fabra, Carrer Dr Aiguader 88, 08003 Barcelona, Spain. Telephone number: +34 93 3160809.

