## Supplementary material for "Effector T cells in poorly perfused tumor regions exhibit a distinct signature of augmented IFN response and reduced PD-1 expression": Riera-Borrull et al Supplemental material

Marta Riera-Borrull *et al*.

**This PDF file includes**:

Table S1
Figures S1 and S2

**Table S1. Primers for RT-qPCR used in this work**

| Primers used for mRNA analysis by real-time quantitative PCR (RT-qPCR) | | |
|---|---|---|
| **Gene product** | **Forward primer (5'-3')** | **Reverse primer (5'-3')** |
| B2m (beta-2 microglobulin) | TTCTGGTGCTTGTCTCACTG | GGAACTGTGTTACGTAGCAG |
| IL-17A | TCAGACTACCTCAACCGTTC | AATTCATGTGGTGGTCCAGC |
| IFNγ | CTCAAGTGGCATAGATGTGG | CAGGTGTGATTCAATGACGC |
| IL-22 | CCTGATGAAGCAGGTGCTAA | TCCTTCAGCCTTCTGACATTCTT |
| L32 | ACCAGTCAGACCGATATGTG | ATTGTGGACCAGGAACTTGC |
| FOXP3 | CAAGGGCTCAGAACTTCTAG | AGCTGATGCATGAAGTGTGG |
| CTLA4 | TCTGAAGCCATACAGGTGACC | TGGTCATTTGTCTGCCGC |
| CXCL10 | AAGGGATCCCTCTCGCAAGGAC | ATCGTGGCAATGATCTCAACAC |
| CCL5 | TTGTCACTCGAAGGAACCGC | AGAGCAAGCAATGACAGGGA |
| STAT1 | TGGTGAAATTGCAAGAGCTG | CAGACTTCCGTTGGTGGATT |
| IFIT1 | TACAGGCTGGAGTGTGCTGAGA | CTCCACTTTCAGAGCCTTCGCA |
| CXCR3 | TGTACCTTGAGGTTAGTGAACG | GGAGTCAGAGAAGTCGCTCT |
| ACSS2 | TCTGCTACAACGTGCTGGAT | CACCCTTCTGAATGCCCTGT |

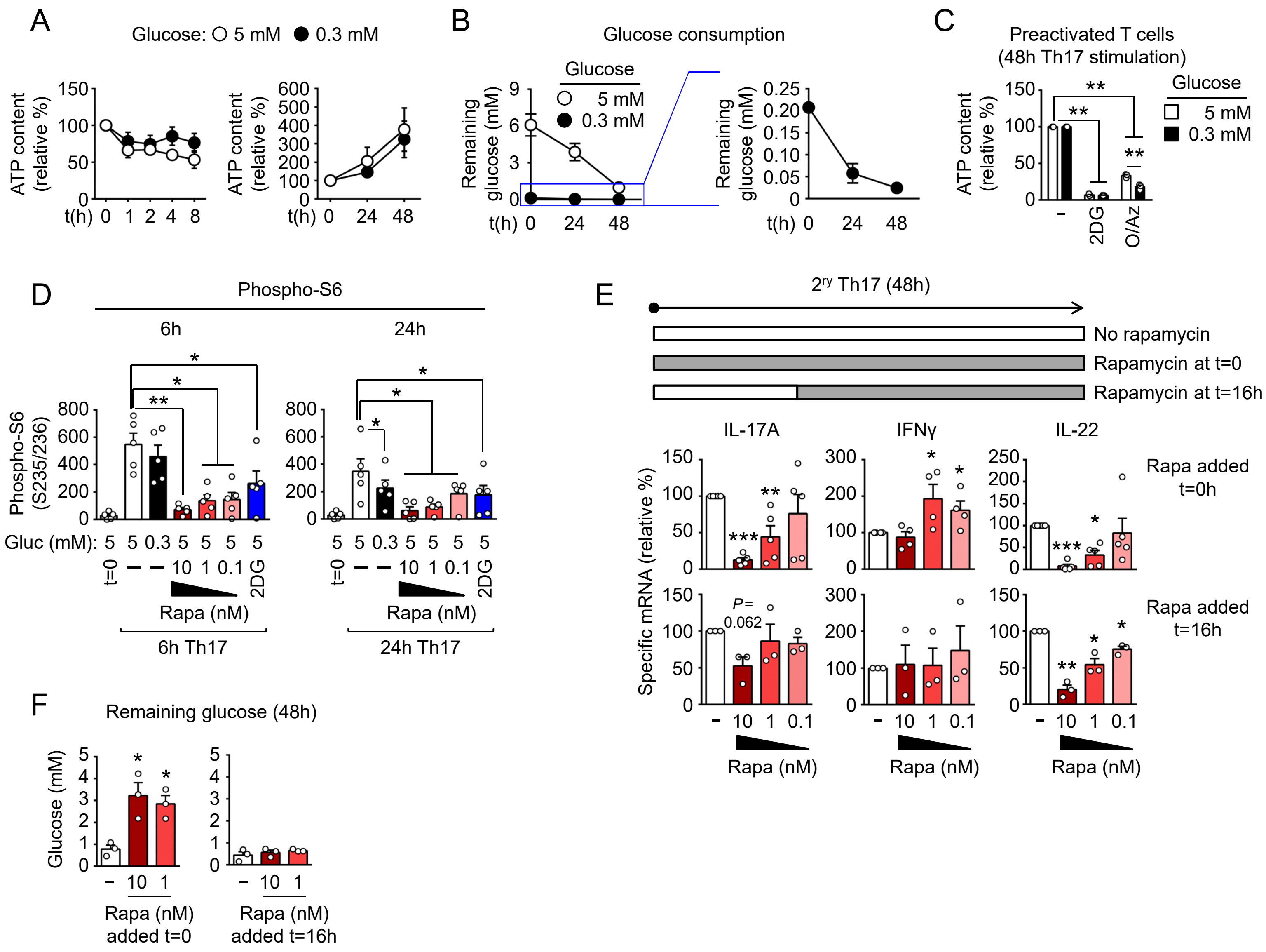


**Figure S1. Activated CD4 T lymphocytes can maintain their glucose-dependent ATP pool and mTORC1 activity under very low glucose levels. (A)** Intracellular ATP content in CD4 cells activated for 5 days in non-polarizing (Th0) conditions and then restimulated as Th17 in 5 or 0.3 mM glucose medium up to 48 hours as in Figure 1A. Results are shown relative to the time of Th17 restimulation (t=0). Results show the mean ± SEM from 3 to 5 independent experiments. Statistical significance was assessed with a one-sample *t* test for comparison with the reference sample at t=0. * $P < 0.05$; ** $P < 0.01$. **(B)** Glucose remaining in the supernatant of cultures of preactivated CD4 cells restimulated as Th17 in 5 or 0.3 mM glucose medium up to 48 hours as in (A). Results show the mean ± SEM from 3 to 4 independent experiments. **(C)** Contribution of glucose and mitochondrial respiration to intracellular ATP levels in CD4 cells after 48 h Th17 stimulation in 5 or 0.3 mM glucose. Inhibitors 2-deoxyglucose (2DG, 5 mM) and oligomycin (0.1 µg/ml) plus sodium azide (20 mM) were added in the last 2 hours of culture. Results show the mean ± SEM from 3 independent experiments. Statistical significance was assessed with a one-sample *t* test for comparison with the reference sample without inhibitors. **(D)** Intracellular levels of phosphorylated S6 in preactivated CD4 cells and then restimulated as Th17 for 6 or 24 hours in 5 or 0.3 mM glucose and with the inhibitors 2DG (2 mM) and rapamycin (0.1 to 10 nM) as indicated. Phospho-S6 was analyzed by flow cytometry and its levels shown as mean fluorescence intensity (MFI). Results show the mean ± SEM from 5 independent experiments. Statistical significance was assessed with a paired *t* test. * $P < 0.05$; ** $P < 0.01$. **(E)** Cytokine mRNA expression in CD4 cells activated for 5 days in non-polarizing (Th0) conditions and then restimulated as Th17 ($2^{ry}$ Th17) in 5 mM glucose for 48 hours, without or with rapamycin added at t=0 or t=16h. Results are shown relative to the time of Th17 restimulation (t=0). Results show the mean ± SEM from 3 to 5 independent experiments. Statistical significance was assessed with a one-sample *t* test for comparison with the reference sample without rapamycin. * $P < 0.05$; ** $P < 0.01$; *P* values below 0.1 are also indicated. **(F)** Glucose remaining in supernatants of cultures of preactivated CD4 cells restimulated as Th17 in 5 mM glucose up to 48 hours, without or with rapamycin as in (E). Results show the mean ± SEM from 3 independent experiments for each rapamycin time point. Statistical significance was assessed with an unpaired *t* test. * $P < 0.05$.

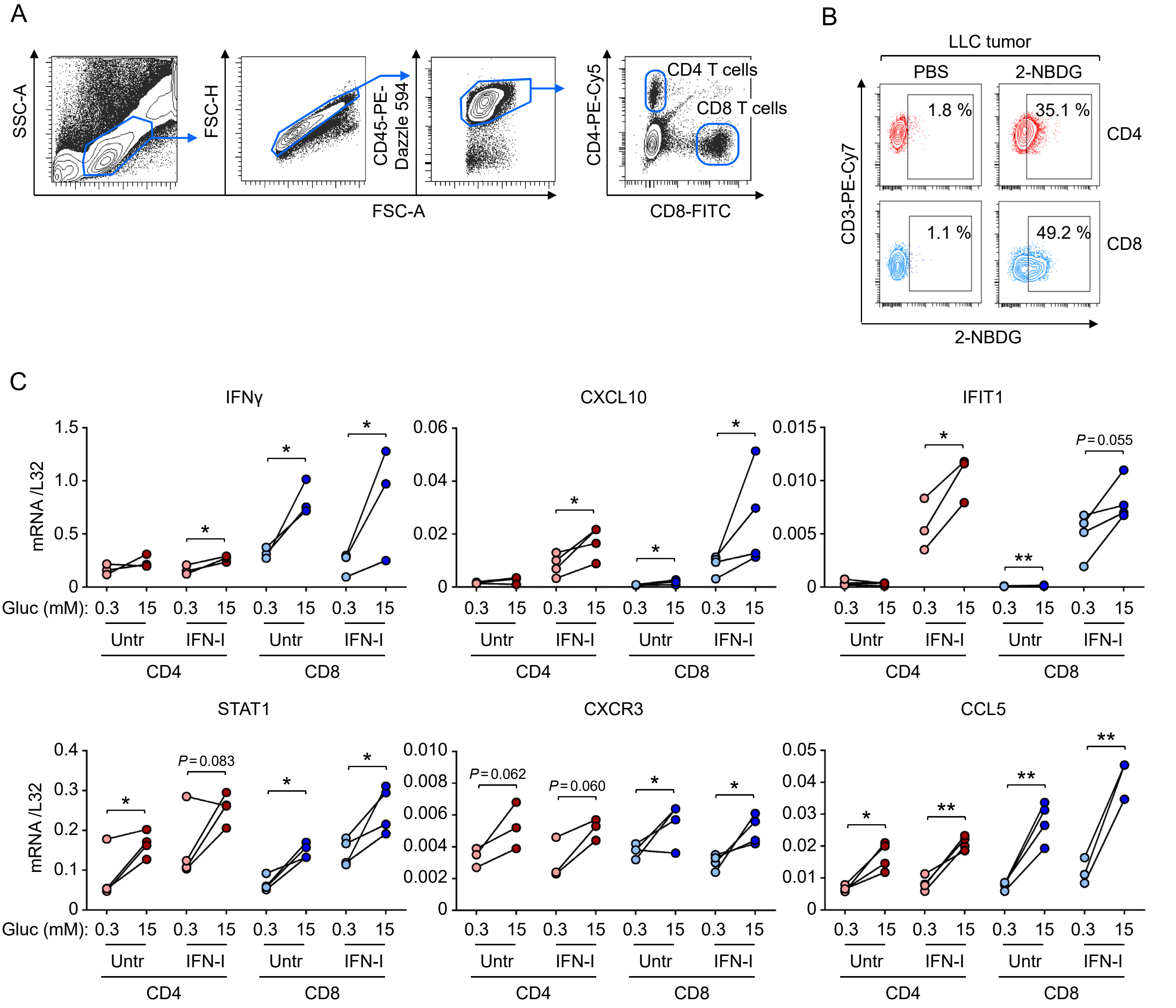


**Figure S2. Tumor-infiltrating CD4 and CD8 T lymphocytes with reduced uptake of 2-NBDG exhibit enhanced expression of glucose-sensitive IFN-I response chemokines and CXCR3 (A)** Gating strategy used to identify CD4 and CD8 T lymphocytes infiltrating LLC tumors. **(B)** A representative experiment illustrating the uptake of 2-NBDG by CD4 and CD8 TILs 15 minutes after having injected the probe intravenously. TILs from PBS-injected mice are shown as negative control. **(C)** Expression of the indicated genes in T lymphocytes preactivated in vitro (5 days) and restimulated (24 hours) without or with a cocktail of IFNα4 and IFNβ1 (IFN-I) in culture medium with 0.3 or 15 mM glucose. T lymphocyte cultures were prepared from 3 to 4 independent mice. Statistical significance was assessed with a paired *t* test. * $P < 0.05$; ** $P < 0.01$. *P* values below 0.1 are also indicated.
